# Mistic: an open-source multiplexed image t-SNE viewer

**DOI:** 10.1101/2021.10.08.463728

**Authors:** Sandhya Prabhakaran, Chandler Gatenbee, Mark Robertson-Tessi, Jeffrey West, Amer A. Beg, Jhanelle Gray, Scott Antonia, Robert A. Gatenby, Alexander R. A. Anderson

## Abstract

Understanding the complex ecology of a tumor tissue and the spatio-temporal relationships between its cellular and microenvironment components is becoming a key component of translational research, especially in immune-oncology. The generation and analysis of multiplexed images from patient samples is of paramount importance to facilitate this understanding. In this work, we present Mistic, an open-source multiplexed image t-SNE viewer that enables the simultaneous viewing of multiple 2D images rendered using multiple layout options to provide an overall visual preview of the entire dataset. In particular, the positions of the images can be taken from t-SNE or UMAP coordinates. This grouped view of all the images further aids an exploratory understanding of the specific expression pattern of a given biomarker or collection of biomarkers across all images, helps to identify images expressing a particular phenotype or to select images for subsequent downstream analysis. Currently there is no freely available tool to generate such image t-SNEs. Mistic is open-source and can be downloaded at: https://github.com/MathOnco/Mistic.

## 1. Introduction

Multiplex imaging of tissues, which allows the simultaneous imaging of multiple biomarkers on a tissue specimen of interest, is a critical tool for clinical cancer diagnosis and prognosis. Historically, patient tissue samples stained with hematoxylin and eosin (H&E) have been used as the gold standard for tumor diagnosis by indicating the presence of tumors and their grade [1] [2] [3]. With the advent of immunohistochemical (IHC) [4] staining and the flourishing of multiplexed imaging approaches that leverage IHC, immunofluorescence (IF), fluorescence *in situ* hybridization (FISH) [5] [6], multiplexed ion beam imaging (MIBI) [7], cyclic labeling (CODEX [8], CyCIF [9]) and imaging mass cytometry (IMC) [10], there is a wealth of potential data to be gleaned from a single section of tissue. Biomarkers can be observed and quantified with their tissue context completely conserved. Due to the multidimensional nature of the data from these multiplexed images, analysis requires computational pipelines to both interrogate and study how the tissue architecture, spatial distribution of multiple cell phenotypes, and co-expression of signaling and cell cycle markers are related and what patterns might exist.

There are several commercial software platforms available for quantifying and analyzing multiplex image data, for example: Imaris (from Oxford Instruments) [11], Amira (from Thermo Fisher Scientific) [12] and Halo (from Indica Labs) [13] [14]. There are also open-source software platforms, for instance: ImageJ [15], CellProfiler [16], V3D [17], BioImageXD [18], Icy [19], FIJI [20], and QuPath [21] for the analysis of two dimensional (2D) biological images. Most of these platforms allow for a single 2D image to be examined at any one time.

A common way to visualize and better understand multidimensional data, such as that coming from multiplex images, is to utilize dimensionality reduction methods such as Uniform Manifold Approximation and Projection (UMAP) [24] or t-distributed Stochastic Neighbor Embedding (t-SNE) [25], where each image is abstracted as a dot in the reduced space. These approaches are especially useful when combined with clustering methods (e.g. Gaussian mixture models [26] [27], Louvain [28], and Leiden [29]) that can highlight key aspects of the data. Whilst utilizing these approaches in our own work dealing with multiplexed images of tumors, we realized that there could be a significant benefit to visualizing the actual tissue samples behind a UMAP or t-SNE scatter projection, thus giving rise to an “image t-SNE”. In our specific application, inspection of the images that constituted each spatially segregated cluster revealed cluster-specific biomarker patterns that, along with the tumor phenotypes, could be mapped succinctly to the therapy response of each patient. Thus, the image t-SNE rendering aided both our understanding and intuition that there exist distinct tumor patterns that guide the clustering, and that these patterns can potentially inform why a specific therapeutic response emerged, leading to further biological insights.

In this work, we present Mistic, an open-source multiplexed image t-SNE viewer that enables the simultaneous viewing of multiple 2D images rendered using multiple layout options to provide an overall visual preview of the entire dataset. In particular, the positions of the images can be taken from t-SNE or UMAP coordinates. This grouped view of all the images further aids an exploratory understanding of the biomarkers’ specific expression pattern across all images, helps to identify images expressing a particular phenotype or to select images for subsequent downstream analysis. Currently there is no freely available tool to generate such image t-SNEs (see Table 1). Software such as BioImageXD and Icy offer do offer a “Gallery” or “Stack Montage” option, where a multichannel image is split into its individual channels to be viewed at once. Mistic is distinct in that *multiple* multichannel images can be processed and rendered at once using either user-predefined coordinates (e.g. from t-SNE or UMAP analysis), random coordinates, or using a grid layout.

**Table 1:**
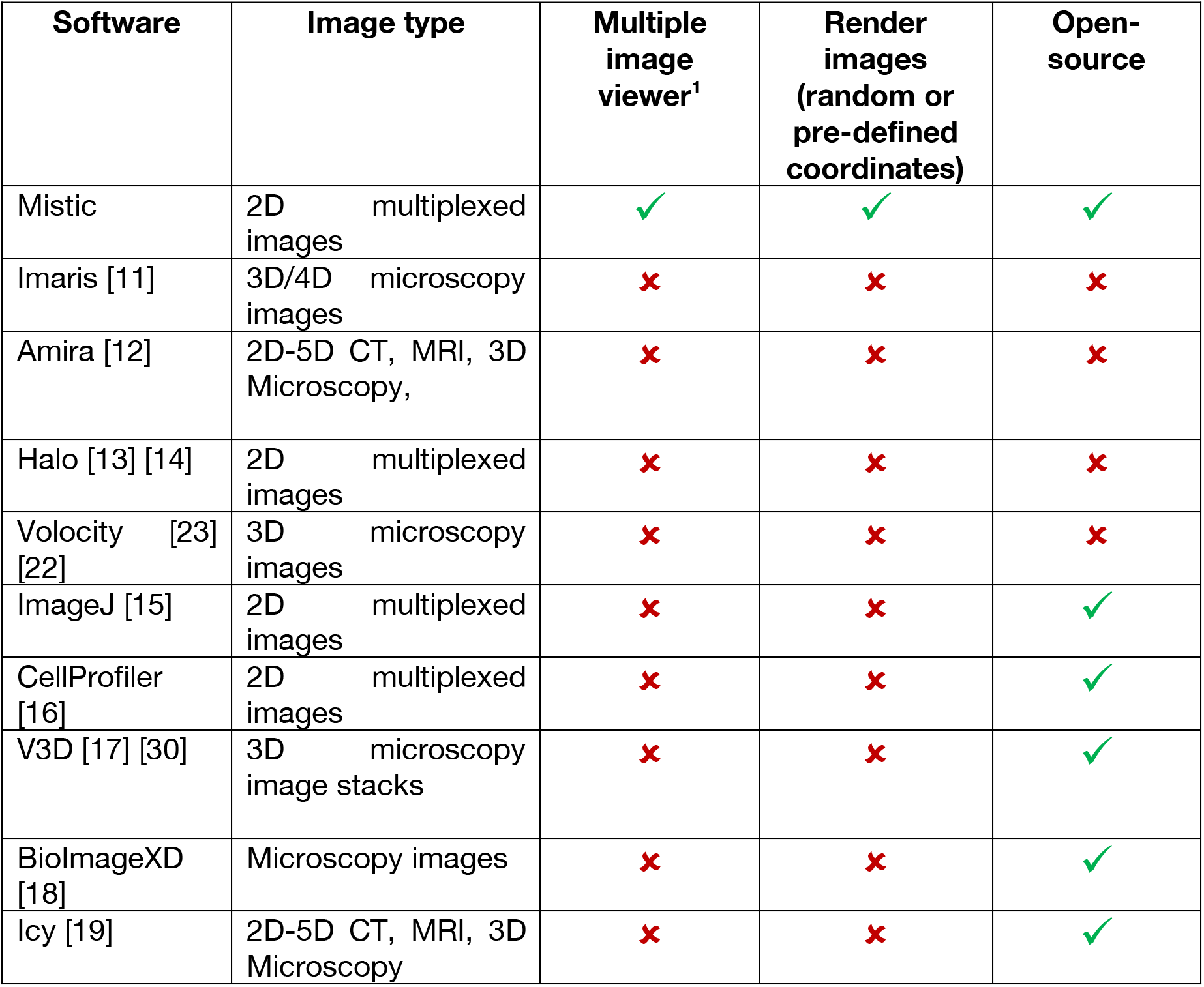
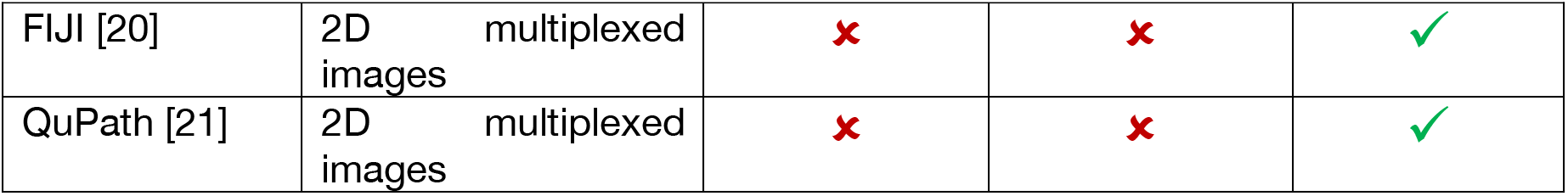
Currently available image analysis software that allow single multiplexed images to be viewed and analyzed, compared with Mistic that allows multiple multiplexed images to be viewed simultaneously, either as an image t-SNE, grid view, or using random coordinates. Mistic has the potential to be integrated into these existing image analysis pipelines as a first step to generate an all-image preview. ^1^ By a multiple image viewer, we mean that each image is a multichannel image itself, unlike a single multichannel image being visualized with its individual channels separated.

| Software | Image type | Multiple image viewer<sup>1</sup> | Render images (random or pre-defined coordinates) | Open-source |
| --- | --- | --- | --- | --- |
| Mistic | 2D multiplexed images | ✓ | ✓ | ✓ |
| Imaris [11] | 3D/4D microscopy images | ✗ | ✗ | ✗ |
| Amira [12] | 2D-5D CT, MRI, 3D Microscopy, | ✗ | ✗ | ✗ |
| Halo [13] [14] | 2D multiplexed images | ✗ | ✗ | ✗ |
| Volocity [23] [22] | 3D microscopy images | ✗ | ✗ | ✗ |
| ImageJ [15] | 2D multiplexed images | ✗ | ✗ | ✓ |
| CellProfiler [16] | 2D multiplexed images | ✗ | ✗ | ✓ |
| V3D [17] [30] | 3D microscopy image stacks | ✗ | ✗ | ✓ |
| BioImageXD [18] | Microscopy images | ✗ | ✗ | ✓ |
| Icy [19] | 2D-5D CT, MRI, 3D Microscopy | ✗ | ✗ | ✓ |
| FIJI [20] | 2D multiplexed images | ✗ | ✗ | ✓ |
| QuPath [21] | 2D multiplexed images | ✗ | ✗ | ✓ |

In Section 2, we illustrate the importance of visualizing multiplexed images using an image t-SNE in the context of non-small cell lung cancer (NSCLC). In Section 3, we describe Mistic and its features, in more detail. We describe code availability in Section 4 and conclude in Section 5.

## 2. Image t-SNE-based visualization of multiplexed images from a NSCLC cohort shows marker expression clustering across different patient response groups

We computationally analyzed 92 7-stain PerkinElmer Vectra images from nine patients with advanced/metastatic non–small cell lung cancer with progression [31]. They were treated with an oral HDAC inhibitor (vorinostat) combined with a PD-1 inhibitor (pembrolizumab). Tumor biopsies were collected from all patients both pre- and on-treatment. Of the nine patients, four qualified as “Response 1” and five as “Response 2”. There are 34 images from patients having Response 1 and 58 images from patients classified as Response 2. Note that we have labeled the clusters, markers, and responses in a generic fashion, since the biological conclusions arising from this data are not the purpose of this work.

We extracted the cell segments per field of view (FoV), built a count matrix with cells as rows and markers as columns, and clustered the count matrix to identify heterogeneous cell types, in particular tumor and immune cells. From these cell types, we automatically demarcate tumor-rich regions, across images.

To further quantify the tumor-immune cell colocalization at the tumor border we cluster the tumor-immune cell counts at the tumor border using a Gaussian mixture model [26] [27]. The clusters are visualized using a standard 2D t-SNE plot where each point represents an image (**Figure 1a, b**). The differentially expressed markers for each of the three clusters are shown in **Figure 1a** and the corresponding patient response of either Response 1 or Response 2 categories, which is known *a priori*, are depicted in **Figure 1b**. We see that there is a higher colocalization of one set of markers for Response 1 patients and higher colocalization of another set of markers for Response 2 patients (**Figure 1a, b**), indicating underlying structural differences between different patient response groups.

**Figure 1:**
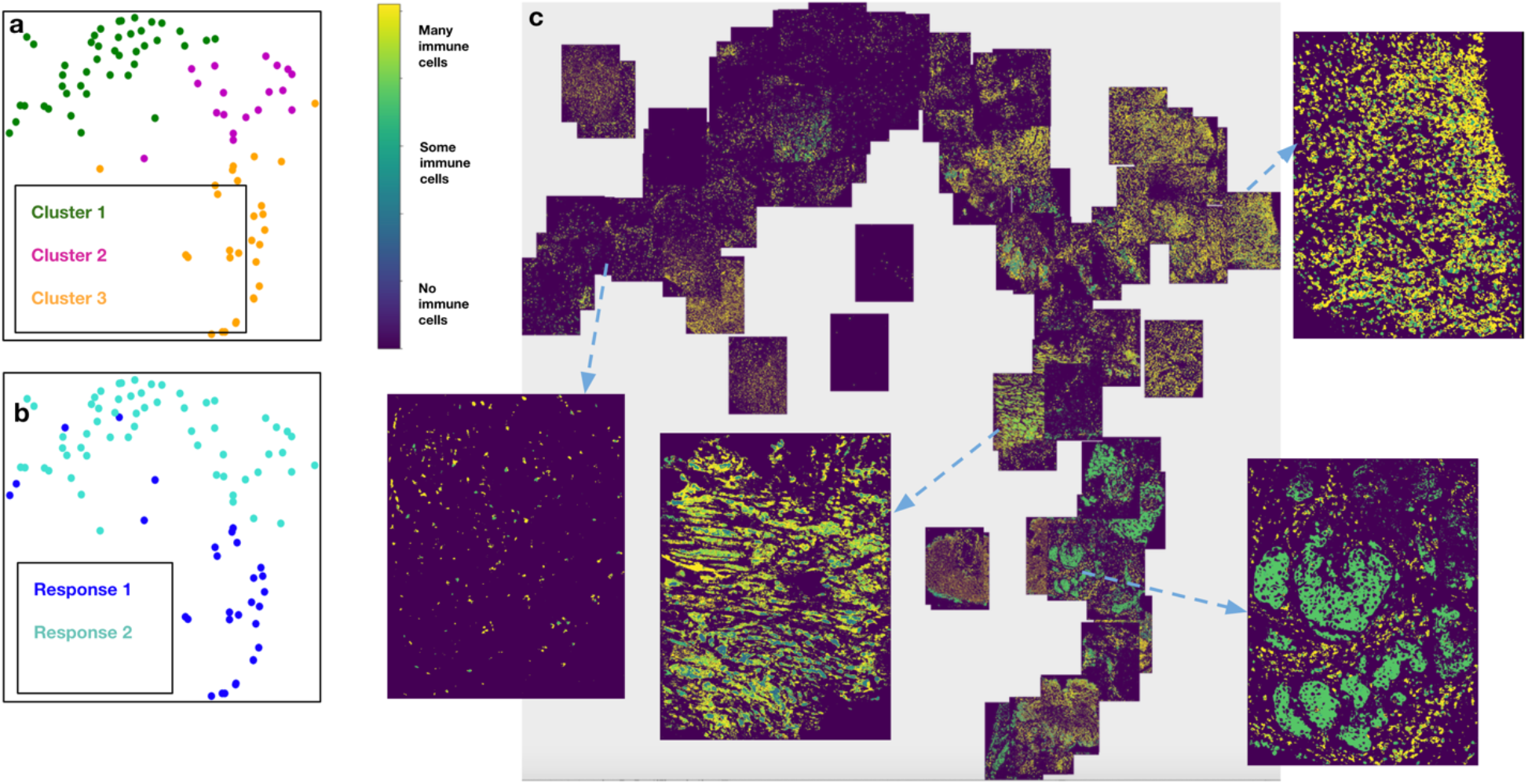
Response 1 and Response 2 patients have significantly different cellular compositions. **a.** 2D t-SNE plot showing three clusters annotated with the differentially expressed markers per cluster. Clusters are obtained using the tumor-immune cell counts at the tumor border where the borders are estimated using convex hulls approximations. **b.** Same t-SNE as in **(a)** depicting the disease response spread. **c.** Image t-SNE using same t-SNE coordinates as in **(a)** illustrating the gradient of immune cells across images. A higher colocalization of immune cells (shown in green) is seen for Response 1 patients.

To better understand how these clusters relate to the actual images, we generated an image t-SNE (**Figure 1c**) where each dot in the t-SNE of Figures 1a and b is replaced with its corresponding multiplexed image. This arrangement of images projected as an image t-SNE clearly highlights the difference in immune cell abundance across Response 1 and Response 2 patient groups.

## 3. Mistic: an open-source multiplexed image t-SNE viewer

In order to facilitate the generation and manipulation of image t-SNEs we developed an image t-SNE viewer called Mistic (multiplexed image t-SNE viewer). Mistic allows the simultaneous viewing of multiple multiplexed images, where images can be arranged using either pre-defined coordinates (e.g. t-SNE or UMAP), randomly generated coordinates, or a grid view. Mistic is written in Python and uses Bokeh [32] which is a Python library for creating interactive visualizations for modern web browsers, along with JavaScript. Mistic has the capability to load and display up to a hundred multiplexed images along with the metadata for the images. It produces publication-ready outputs that can be saved in PNG format. Additionally, it can be used as the initial image viewer for exploratory image analysis before switching to more comprehensive (but single-image) viewers such as ImageJ [15], Fiji [20], and QuPath [21].

Mistic provides many of the standard image-viewing features that users have come to rely on and expect, through a user-input panel and two canvases. The user-input panel (**Figure 2a**) allows the user to select: markers for rendering the multiplexed images; optional image borders; the arrangement of the images by coordinates or grid; and the option to shuffle the order of image rendering for overlapping images. An overall color theme for Mistic can be chosen from black, blue and gray. Mistic further provides two canvases for image t-SNE rendering: a static canvas showing the image t-SNE with all the multiplexed images (**Figure 2b**), which is generated based on user preferences; and a live canvas depicting the corresponding t-SNE scatter plot that uses the metadata from the images, where each image is represented as a dot (**Figure 2c**). We explain the two canvases in detail in the following subsections (3.1 and 3.2).

**Figure 2:**
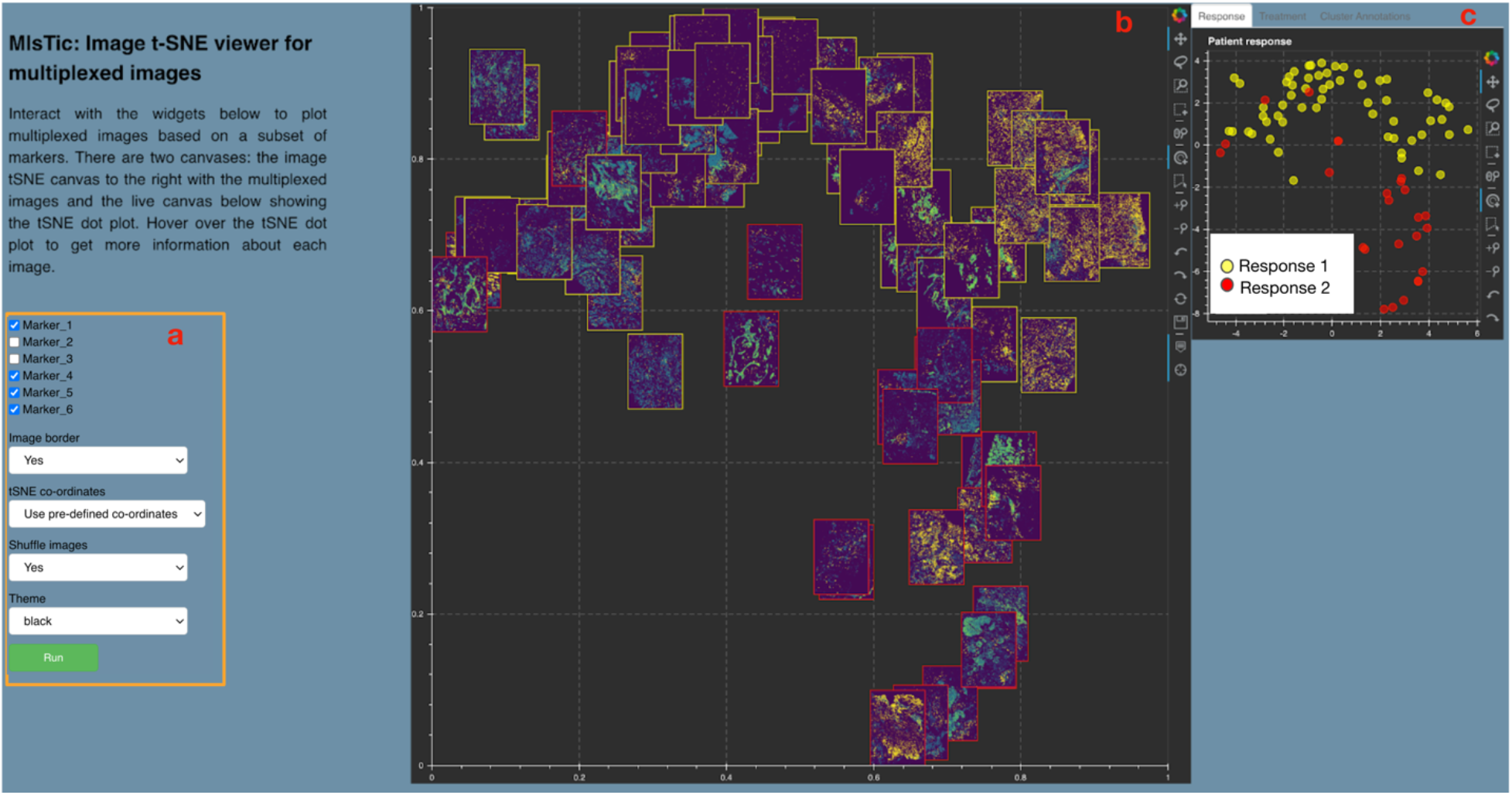
Mistic GUI. **a.** User-input panel where markers can be selected, images borders can be added, new or pre-defined image display coordinates can be chosen, and a theme for the canvases can be selected. **b.** Static canvas showing the image t-SNE colored and arranged as per user inputs. **c.** Live canvas showing the corresponding t-SNE scatter plot where each image is represented as a dot. The live canvas has tabs for displaying additional information per image. Metadata for each image can be obtained by hovering over each dot.

### 3.1 Image t-SNE rendered through the static canvas

To view the multiplexed images simultaneously, Mistic offers the user the ability to choose from three different image layouts (see **Figure 3**): (i) t-SNE layout based on user-predefined coordinates; (ii) vertical grid arrangement of all images; (iii) random layout based on coordinates that Mistic generates. Depending on the specific layout chosen, the live canvas will be updated accordingly (see section 3.2). The user can also opt to shuffle the front-to-back order in which images are rendered, as shown in **Figure 4**; this is particularly useful when there are many overlapping images. These options can be chosen from the user-input panel (**Figure 2a**).

**Figure 3:**
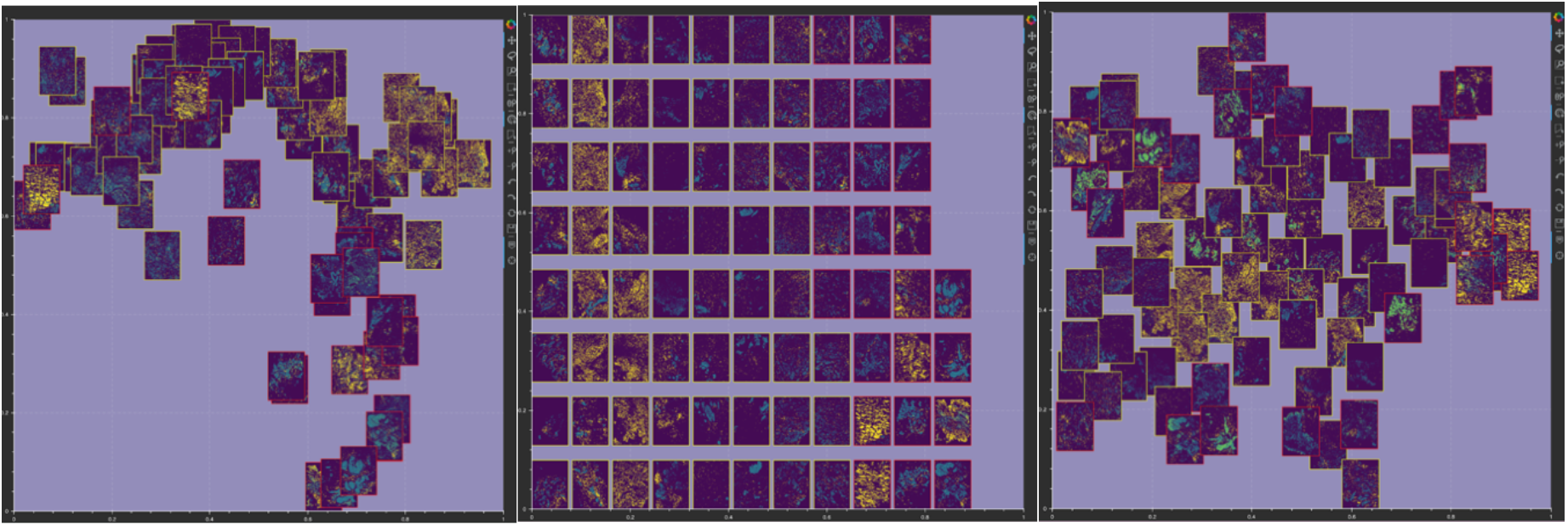
Image layout in Mistic’s static canvas: (i) based on user-defined t-SNE coordinates (left); (ii) vertical rows (center); (iii) randomly placed (right).

**Figure 4:**
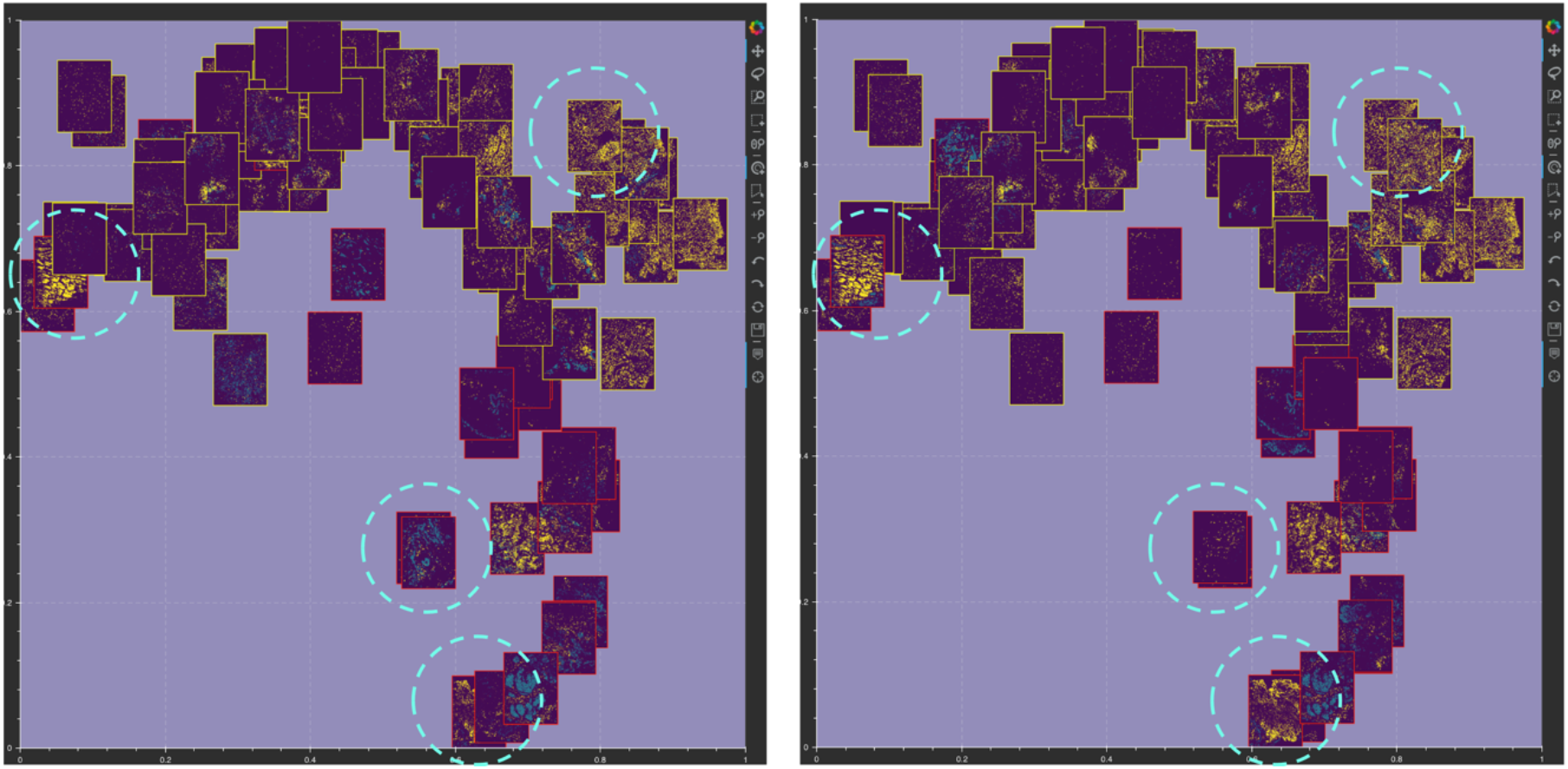
Mistic can shuffle the order in which images are rendered on the static canvas. Examples of images being shuffled between two renderings are marked in dotted circles.

Additional options available to users include: (i) choosing the canvas color theme (black, gray or dark blue), and (ii) applying borders around images to highlight specific metadata about the images. In **Figure 2b**, for the example from our NSCLC cohort, the borders indicate images belonging to either Response 1 (yellow border) or Response 2 (red border) patients.

### 3.2 Metadata rendered through the live canvas

The live canvas of Mistic offers different metadata renderings of the multiplexed images through t-SNE scatterplots where every multiplexed image is a dot on the scatterplot. In our NSCLC example, Mistic renders the t-SNE scatter plot based on either,

i. Response category of the patients (e.g. based on RECIST classification),
ii. Treatment phase (such as pre-treatment or during treatment),
iii. Cluster annotations that are based on the differential-expression analysis of the markers. This metadata information must be provided by the user, using appropriate folders provided in Mistic’s code repository, available here: https://github.com/MathOnco/Mistic.

**Figure 5:**
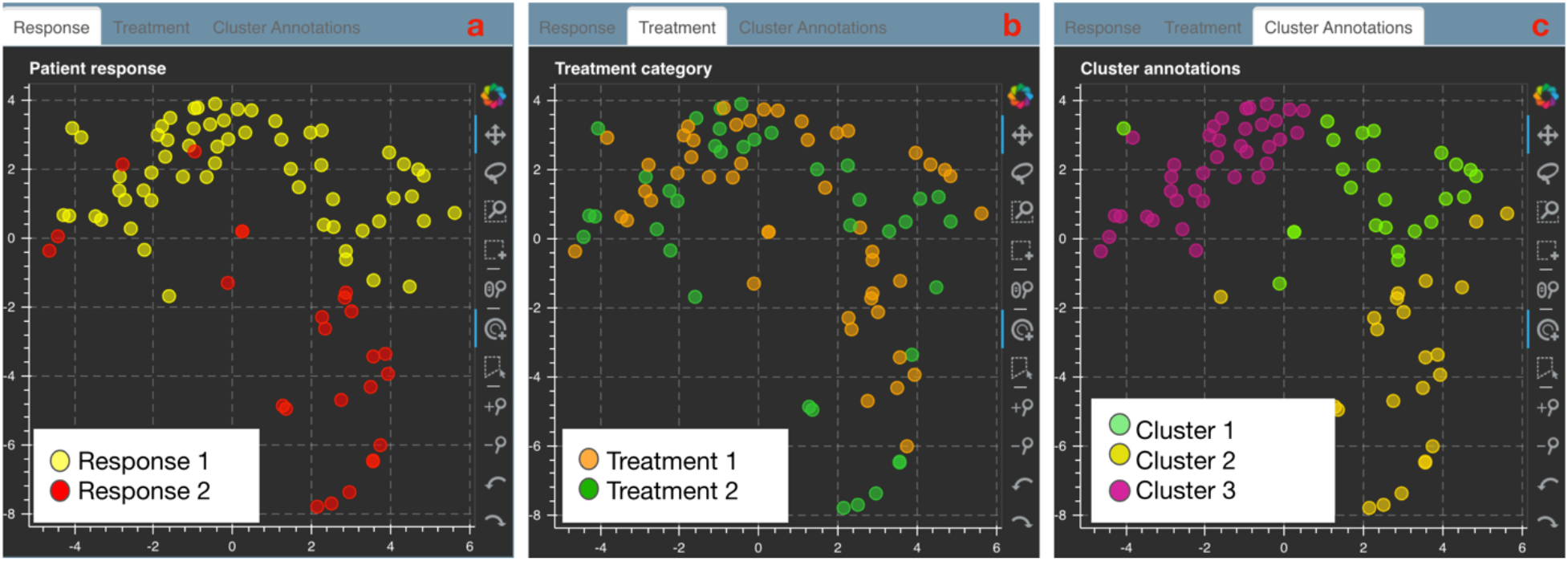
Live canvas of Mistic rendering the t-SNE scatter plot with different image metadata for our NSCLC example. The metadata consists of response category (panel a), treatment phase (panel b) and cluster annotations based on marker expression (panel c).

### 3.3 Processing user inputs from Mistic GUI

#### a. Image processing based on markers selected

Each user-selected marker channel of the multiplexed image is denoised separately using Otsu thresholding and combined to create a denoised image. This image is stored as an array in the unsigned byte format (‘uint8’) to enable easy format conversion.

#### b. Border option

An image with a border is created by pasting the cleaned image onto a rectangle with a slightly larger height and width than the cleaned image. The rectangle is filled with a color based on the metadata provided by the user.

#### c. Thumbnail generation

A thumbnail is a concise representation of the original multiplexed image rendered based on user selections. Thumbnails are created for all multiplexed images by downscaling and resizing the height and width of the cleaned image, and are also saved in the user output folder.

#### d. Random co-ordinate creation

To generate a set of non-clustered random sample of 2D points, we use a modified version of the ‘Poisson disc sampling’ approach [33] [34].

Finally, all the thumbnails are pasted onto a larger 2D image that gets rendered onto Mistic’s static canvas where the thumbnails are positioned based on pre-defined coordinates (e.g. t-SNE or UMAP), randomly generated coordinates, or as vertical grids.

### 3.4 Bokeh plot tools

Each Mistic canvas uses the interactive Bokeh toolbar to save plots, select regions, and change plot parameters such as zoom level, reset, pan, etc. **Figure 6** shows the set of plot tools used. Further documentation of the Bokeh toolbar and how to use it can be found here: https://docs.bokeh.org/en/latest/docs/user_guide/tools.html

**Figure 6:**
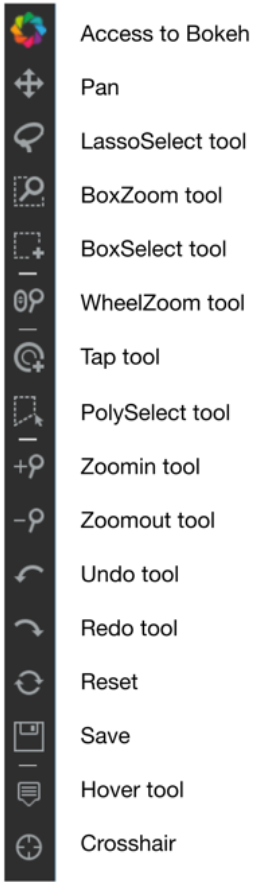
Interactive Bokeh plot tools used in Mistic.

## 4. Code availability

Mistic is downloadable at: https://github.com/MathOnco/Mistic. Instructions regarding installation, setup and code deployment can be found there. The code is written in Python 3.6 and uses Bokeh which is a Python library for creating interactive visualizations for modern web browsers. Mistic has been tested using 92 7-channel Vectra images each of 1024×1024 pixels in dimension.

## 5. Conclusion

Understanding the complex ecology of a tumor tissue and the spatio-temporal relationships between its cellular and microenvironment components is becoming a key component of translational research, especially in immune-oncology. The generation and analysis of multiplexed images from patient samples is of paramount importance to facilitate this understanding. In Table 1, we highlight different image viewers currently available as open-source or commercial software. To our knowledge, there exists no current image viewer allowing the simultaneous preview of multiple multiplexed images, rendered using t-SNE coordinates or random coordinates. Mistic aims to fill this gap by providing this simple functionality to view multiple images at once, whilst also giving users the option to view images based on a set of user choices. In our test runs using 92 images (with dimension 1024×1024 pixels), Mistic takes under a minute to process and render the images according to the user options available (for user options, see **Section 3.1** and **Figure 2a**).

As part of future work, a few potential improvements will be introduced to Mistic. Once a set of images are identified using Mistic, we would like to render those images separately in the live panel. This gives the user an additional perspective to refine the selected images for further analysis. We also hope to integrate Mistic into one of the open-source software viewers as listed in Table 1. This would require the development of an additional framework in React JavaScript [35], which is the single largest User Interface framework.

Through our generalized example of NSCLC consisting of 92 images from nine patients, we have demonstrated the functionality and practicality of Mistic. Our aim is that Mistic will be used as a first step to viewing multiplexed images simultaneously. This all-image visual preview should facilitate preliminary insights into possible marker expression patterns, aiding downstream image analysis for predicting disease progression and identifying clinical biomarkers.

## 7. Contributions

Conceptualization, S.P., and A.R.A.A.; Manuscript preparation, S.P., and A.R.A.A; Manuscript editing, S.P., C.G., M.R.-T., A.B., J.G., S.A., R.A.G., and A.R.A.A; Software design and implementation, S.P.; Software testing, S.P., and J.W.; Supervision, S.A., R.A.G., and A.R.A.A; Funding acquisition, S.A., R.A.G., and A.R.A.A;

## 8. Acknowledgments

The authors gratefully acknowledge funding by the National Cancer Institute via the Cancer Systems Biology Consortium (CSBC) U01CA232382 (supporting S.P., C.G., M.R.-T., A.R.A.A), the Physical Sciences Oncology Network (PSON) U54CA193489 (supporting C.G., M.R.-T., J.W., A.R.A.A), and support from the Moffitt Center of Excellence for Evolutionary Therapy (supporting M.R.-T., and A.R.A.A).

## Bibliography

[1] R. Dapson and R. Horobin, “Dyes from a twenty-first century perspective,” Biotechnic & Histochemistr, vol. 84, no. 4, pp. 135–137, 2009.

[2] M. Titford, “The long history of hematoxylin,” Biotechnic & Histochemistry, vol. 80, no. 2, pp. 73–80, 2005.

[3] Z. Saadatpour, A. Rezaei, H. Ebrahimnejad, B. Baghaei, G. Bjorklund, M. Chartrand, A. Sahebkar, H. Morovati, H. R. Mirzaei and H. Mirzaei, “Imaging techniques: new avenues in cancer gene and cell therapy,” Cancer Gene Therapy, vol. 24, pp. 1–5, 2017.

[4] W. Tan, S. N. Nerurkar, H. Y. Cai, H. Ng, D. Wu, Y. Wee, J. Lim, J. Yeong and T. Lim, “Overview of multiplex immunohistochemistry/immunofluorescence techniques in the era of cancer immunotherapy,” Cancer communications (London, England), vol. 40, no. 4, pp. 135–153, 2020.

[5] J. Moffitt, D. Bambah-Mukku, S. Eichhorn, E. Vaughn, K. Shekhar, J. Perez, N. Rubinstein, J. Hao, A. Regev, C. Dulac and X. Zhuang, “Molecular, spatial, and functional single-cell profiling of the hypothalamic preoptic region,” Science, vol. 362, no. 6416, 2018.

[6] N. Ji and A. Van Oudenaarden, “Single molecule fluorescent in situ hybridization (smFISH) of C. elegans worms and embryos,” WormBook, 2012.

[7] L. Keren, M. Bosse, S. Thompson, T. Risom, K. Vijayaragavan, E. McCaffrey, D. Marquez, R. Angoshtari, N. Greenwald, H. Fienberg and J. Wang, “MIBI-TOF: A multiplexed imaging platform relates cellular phenotypes and tissue structure,” Science advances, vol. 5, no. 10, 2019.

[8] Y. Goltsev, N. Samusik, J. Kennedy-Darling, S. Bhate, M. Hale, G. Vazquez, S. Black and G. P. Nolan, “Deep profiling of mouse splenic architecture with CODEX multiplexed imaging,” Cell, vol. 174, no. 4, pp. 968–981, 2018.

[9] J. Lin, B. Izar, S. Wang, C. Yapp, S. Mei, P. Shah, S. Santagata and P. Sorger, “Highly multiplexed immunofluorescence imaging of human tissues and tumors using t-CyCIF and conventional optical microscopes,” Elife, vol. 7, 2018.

[10] C. Giesen, H. A. Wang, D. Schapiro, N. Zivanovic, A. Jacobs, B. Hattendorf, P. Schüffler, D. Grolimund, J. Buhmann, S. Brandt, Z. Varga and B. Bodenmiller, “Highly multiplexed imaging of tumor tissues with subcellular resolution by mass cytometry.,” Nature Methods, vol. 11, no. 4, pp. 417–422, 2014.

[11] I. Oxford, “Imaris Microscopy Image Analysis,” [Online]. Available: https://imaris.oxinst.com/packages.

[12] T. Scientific, “Amira for Life & Biomedical Sciences,” [Online]. Available: https://www.thermofisher.com/us/en/home/industrial/electron-microscopy/electron-microscopy-instruments-workflow-solutions/3d-visualization-analysis-software/amira-life-sciences-biomedical.html.

[13] Indica Lab, “Indica Lab,” [Online]. Available: https://indicalab.com/.

[14] Labs, Indica, “Halo 3.1 Tissue Classifier Ste-by-step guide,” April 2020. [Online]. Available: https://learn.indicalab.com/wp-content/uploads/2020/08/Indica-Labs-Tissue-Classifier-1.pdf. [Accessed May 2021].

[15] M. Abramoff, P. Magalhaes and S. Ram, “Image processing with ImageJ,” Biophotonics International, vol. 11, pp. 36–42, 2004.

[16] A. Carpenter, “CellProfiler: image analysis software for identifying and quantifying cell phenotypes,” Genome Biology, vol. 7, no. R100, 2006.

[17] H. Peng, Z. Ruan, F. Long, J. Simpson and E. Myers, “V3D enables real-time 3D visualization and quantitative analysis of large-scale biological image data sets,” Nature Biotechnology, vol. 28, pp. 348–353, 2010.

[18] P. Kankaanpää, L. Paavolainen, S. Tiitta, M. Karjalainen, J. Päivärinne, J. Nieminen, V. Marjomäki, J. Heino and D. J. White, “BioImageXD: an open, general-purpose and high-throughput image-processing platform,” Nature Methods, vol. 9, no. 7, pp. 683–689, 2012.

[19] F. de Chaumont, S. Dallongeville and Olivo-Marin, “ICY: a new opensource community image processing software,” IEEE Int. Symp. on Biomedical Imaging, pp. 234–237, 2011.

[20] J. Schindelin, I. Arganda-Carreras and E. Frise, “Fiji: an open-source platform for biological-image analysis,” Nature Methods, vol. 9, p. 676–682, 2012.

[21] P. Bankhead, “QuPath: Open source software for digital pathology image analysis,” Scientific Reports, 2017.

[22] Elmer, Perkin, “Volocity User Guide,” [Online]. Available: http://physiology.med.unc.edu/Resources/Volocity/VolocityUserGuide60.pdf.

[23] P. Elmer, “Volocity 3D Image Analysis Software,” [Online]. Available: https://resources.perkinelmer.com/lab-solutions/resources/docs/BRO_VolocityBrochure_PerkinElmer.pdf?_ga=2.16883582.613118504.1621957415-1084019245.1621957415. [Accessed May 2021].

[24] E. Becht, L. McInnes, J. Healy, C. Dutertre, I. Kwok, L. Ng, F. Ginhoux and E. Newell, “Dimensionality reduction for visualizing single-cell data using UMAP,” Nature biotechnology, vol. 37, no. 1, pp. 38–44, 2019.

[25] L. Van Der Maaten and G. Hinton, “Visualizing high-dimensional data using t-SNE,” Journal of Machine Learning Research, vol. 9, no. 26, 2008.

[26] G. J. McLachlan and K. E. Basford, “Mixture models: Inference and applications to clustering,” 1988.

[27] A. Dempster, N. Laird and D. Rubin, “Maximum likelihood from incomplete data via the EM algorithm,” Journal of the Royal Statistics Society, vol. 39, no. 1, pp. 1–38, 1977.

[28] V. D. Blondel, J.-L. Guillaume, R. Lambiotte and E. Lefebvre, “Fast unfolding of communities in large networks,” J. Stat. Mech. Theory Exp, vol. 10008, no. 6, 2008.

[29] V. Traag, L. Waltman and N. van Eck, “From Louvain to Leiden: guaranteeing well-connected communities,” Science Reports, vol. 9, no. 5233, 2019.

[30] H. Peng, “V3D User Manual,” 2009. [Online]. Available: https://www.cgl.ucsf.edu/chimera/data/v3d-mar2010/V3D_Manual_v2.031_simple.pdf.

[31] J. E. Gray, A. Saltos, T. Tanvetyanon, E. B. Haura, B. Creelan, S. J. Antonia, M. Shafique, H. Zheng, W. Dai, J. J. Saller, Z. Chen, N. Tchekmedyian, K. Goas, R. Thapa, T. A. Boyle and Chen, “Phase I/Ib Study of Pembrolizumab Plus Vorinostat in Advanced/Metastatic Non–Small Cell Lung Cancer,” Clinical Cancer Research, vol. 25, no. 22, pp. 6623–6632, 2019.

[32] Bokeh Development Team, “Bokeh: Python library for interactive visualization,” 2018. [Online]. Available: http://www.bokeh.pydata.org.

[33] R. Bridson, “Fast Poisson Disk Sampling in Arbitrary Dimensions,” ACM SIGGRAPH sketches, 2007.

[34] C. Hill, Learning Scientific Programming with Python, Cambridge University Press, 2020, p. 568.

[35] “React - A JavaScript library for building user interfaces,” [Online]. Available: https://reactjs.org/.

[36] M. Berthold, “KNIME: the Konstanz Information Miner in Studies in Classification, Data Analysis, and Knowledge Organization,” Springer, pp. 319–326, 2007.

[37] R. Rashid, Y. Chen, J. Hoffer, J. Muhlich, J. Lin, R. Krueger, H. Pfister, R. Mitchell, S. Santagata and P. Sorger, “Interpretative guides for interacting with tissue atlas and digital pathology data using the Minerva browser,” BioRxiv, 2020.

[38] J. Hoffer, R. Rashid, J. L. Muhlich, Y. Chen, D. Russell, J. Ruokonen, R. Krueger, H. Pfister, S. Santagata and S. Pk, “Minerva: a light-weight, narrative image browser for multiplexed tissue images.,” Journal of Open Source Software, vol. 5, no. 54, p. 2579, 2020.

